# Asymmetric directed functional connectivity within the frontoparietal motor network during motor imagery and execution

**DOI:** 10.1101/2020.10.22.351106

**Authors:** Takeshi Ogawa, Hideki Shimobayashi, Jun-ichiro Hirayama, Motoaki Kawanabe

## Abstract

Both imagery and execution of motor controls consist of interactions within a neuronal network, including frontal motor-related regions and posterior parietal regions. To reveal neural representation in the frontoparietal motor network, two approaches have been proposed thus far: one is decoding of actions/modes related to motor control from the spatial pattern of brain activity; another is to estimate directed functional connectivity, which means a directed association between two brain regions within motor regions. However, directed connectivity among multiple regions of the motor network during motor imagery (MI) or motor execution (ME) has not been investigated. Here, we attempted to characterize the directed functional connectivity within the frontoparietal motor-related networks between the MI and ME conditions. We developed a delayed sequential movement and imagery task to evoke brain activity associated with data under ME and MI via functional magnetic resonance imaging scanning. We applied a causal discovery approach, linear non-Gaussian acyclic causal model, to identify directed functional connectivity among the frontoparietal motor-related brain regions for each condition. We demonstrated higher directed functional connectivity from the contralateral dorsal premotor cortex (dPMC) to the primary motor cortex (M1) in ME than in MI. We mainly identified significant direct effects of the dPMC and ventral premotor cortex (vPMC) to the parietal regions. In particular, connectivity from the dPMC to the superior parietal lobule in the same hemisphere showed significant positive effects across all conditions. Contrastingly, interlateral connectivities from the vPMC to the superior parietal lobule showed significantly negative effects across all conditions. Finally, we found positive effects from A1 to M1 in the same hemisphere, such as the audio-motor pathway. These results indicated that the sources of motor command originated in d/vPMC influenced M1 and parietal regions as achieving ME and MI. Additionally, sequential sounds may functionally facilitate temporal motor processes.

## 1. Introduction

Achievements of motor imagery (MI) and motor execution (ME) consist of interactions within a neuronal network, including frontal motor regions and posterior parietal regions. According to previous studies, MI is defined as a mental simulation or a mental rehearsal of movements without any overt physical movements (Hanakawa et al., 2008; Lorey et al., 2014; Pilgramm et al., 2016). Neural activities during MI can be predominantly modulated by tasks containing visual, auditory, and kinesthetic aspects (Hanakawa, 2016). Neural substrates of the motor system have common and specific representations corresponding to MI or ME.

Previous studies have reported that information associated with MI can be extracted from activation patterns represented in the frontoparietal network (Apšvalka et al., 2018; Gallivan et al., 2011, 2013; Gao et al., 2011; Hanakawa et al., 2008; Kasess et al., 2008; Lorey et al., 2014; Nambu et al., 2015; Pilgramm et al., 2016; Yokoi & Diedrichsen, 2019; Zabicki et al., 2017). Anatomical brain regions in the frontoparietal network are associated with the following motor processes: i) the primary motor cortex (M1) and primary somatosensory cortex (S1) are activated during sequential movement; ii) the dorsal premotor cortex (dPMC), ventral premotor cortex (vPMC), and supplementary motor area (SMA) are activated during the preparation period of the movement (Hanakawa et al., 2008; Nambu et al., 2015); iii) neural activation in parietal regions, including the inferior parietal sulcus (IPS), inferior parietal lobule (IPL), and superior parietal lobule (SPL) represents gestures (Pilgramm et al., 2016; Zabicki et al., 2017) and target locations depending on head- or eye-centered coordinates and arm/limb-centered coordinates (Gallivan et al., 2011). These findings suggest that MI and ME are achieved by interactive communications between brain regions in the frontoparietal network.

To evaluate the functional relationship between multiple brain regions associated with MI and ME, it is important to investigate the directed functional connectivity of the frontoparietal network, including both hemispheres (Hanakawa, 2016). Previously, several studies have identified the directed functional connectivity of ME and MI, such as the relationship between SMA and M1 using dynamic causal modeling (DCM) (Kasess et al., 2008; Penny et al., 2004) and a frontoparietal network using Granger causality mapping (GCM) (Deshpande et al., 2008; Gao et al., 2011). DCM is one of the approaches to estimate directed functional connectivity based on a neurophysiological model. However, selecting the best model (i.e., connectivity pattern) from all possible ones is often infeasible in DCM, especially, given many brain regions and related factors of interest (Smith et al., 2011). GCM is a data-driven method and can be more efficient than DCM. Its variant named conditional GCM (CGCM: Ding et al., 2006) was applied to analyzing fMRI data of ME/MI conditions (Gao et al., 2011; Wang et al., 2019), demonstrating an asymmetry of directed functional connectivity in right-handed subjects. However, GCM/CGCM for many brain regions is not very straightforward as it typically needs to evaluate the Granger causality measure separately for every single connection.

In the frontoparietal network, asymmetry between brain regions contralateral and ipsilateral to a manipulating hand is a characteristic aspect of motor-related processing that arises depending on handedness. In general, M1 in both hemispheres is activated during MI and ME, but the region contralateral to a controlled hand shows relatively higher activity than the ipsilateral region because of inhibition from the contralateral M1 to the one ipsilateral to the hand in adults (Hanakawa et al., 2005). Ipsilateral M1, however, also encodes single finger movements during ME, even though the mean activation is lower as revealed by a decoding analysis (Diedrichsen et al., 2013). The asymmetric properties of motor control can appear not only in the mean activation or decoding accuracies, but also in directed functional connectivity. Using conditional GCM, bidirectional edges of directed functional connectivities were extracted in the left-hand task among L-SPL, L-IPL, and L-dPMC; therefore, selected edges for the right-hand and left-hand tasks were not the same (Gao et al., 2011). However, asymmetric network structures of the various motor-associated regions in the frontoparietal network and related networks, in particular with both hemispheres, have been poorly understood.

Here, we investigated the asymmetric network structure related with moto-associated regions represented by the directed functional connectivity during MI and ME. To investigate the network structure, we developed a delayed sequential movement and imagery (dSMI) task by extending the one proposed by Hanakawa et al. (2008) to acquire enough fMRI data samples under ME and MI. We applied a novel causal discovery approach, DirectLiNGAM (Shimizu et al., 2011) with the fMRI time courses in the motor-related regions of interest (ROIs) in order to estimate their directed functional connectivity for each condition. DirectLiNGAM is an algorithm to extract causal links based on a linear non-Gaussian acyclic causal model (LiNGAM: Shimizu et al., 2006). It is particularly suitable to analyze large-scale connectivity structures with many ROIs, as compared to DCM/GCM. Finally, we statistically evaluated the directed functional connectivity within frontal regions, between frontoparietal regions, and between audio-motor regions, and illustrated the symmetric/asymmetric structure of the network.

## 2. Methods

### 2.1 Participants

A total of 27 adults (3 female, mean age = 25.4 ± 5.9 years) participated in this study. Participants received cash remuneration for their involvement in this experiment. All participants had normal or corrected-to-normal vision, no reported history of drugs that affected the central nervous system, and no neurological disease. All participants were right-handed, as assessed by the Edinburgh Handedness Inventory (Oldfield, 1971). All participants provided written informed consent prior to the experiment. This study was approved by the ethical committee of the Advanced Telecommunications Research Institute International and followed the Declaration of Helsinki.

### 2.2 Delayed sequential movement and imagery task

We conducted a dSMI task, which was modified from the original version (Hanakawa et al., 2008) because we needed to collect enough trials within a limited fMRI experiment time. Because of our modification, we collected data samples that were sufficient to optimize the directed functional connectivity with a causal discovery approach. We wrote an in-house program coded for the dSMI task using Psychophysics Toolbox extensions (Brainard, 1997; Kleiner et al., 2007; Pelli, 1997) in MATLAB (MathWorks Inc., Natick, USA). Briefly, each run incorporated both performance modes (ME, MI, and PL: passive listening) in a pseudorandom manner. Three different classes of visual stimuli (Move: ME; Image: MI; Listen: PL) were alternately presented (Fig. 1). As an instruction stimulus (IS), an Arabic numeral 1, 2, or 3 was visually presented indicate the initial finger of tapping sequence (1 = the pointing finger, 2 = the middle finger, 3 = the ring finger). The initial finger was semi-randomly selected at each trial. After the IS, a cue stimulus (CS) was presented for 2 s. Each participant was instructed to sequentially tap following short beep sounds (Move or Image) or passively listening to the beep sounds (Listen) in one of the modes, ME, MI, and PL, respectively. The participants were required to press the response buttons corresponding to the beep sound (duration = 0.2 s; tone = 500 Hz; frequency = 0.667 Hz) following the instructed sequence in response to the Move stimulus (Move trials) during the induction period (IP), and press the response button corresponding to the next finger. In response to the Image stimulus, the participants were asked to image the finger tapping during the IP (7 s) from the instructed finger as performed by him/her (first-person perspective) without actual finger tapping (Image trials), and then, were required to press the response button that corresponds to the next finger. In the case of Listen, the participants were asked to respond not only without the movement of their fingers but also without imagining their finger movements during the IP and answer periods. During the MI and PL conditions, we confirmed the absence of finger movement in the absence of the button’s response. The Move, Image, and Listen stimuli were pseudo-randomly assigned to the cue stimuli; therefore, the participants were unable to predict the forthcoming performance mode. The interstimulus interval (ISI) pseudo-randomly lasted for 8, 10, or 12 s. We did not analyze the PL condition in further analysis. Finally, we collected 30 trials in four conditions (right ME: R-ME; right MI: R-MI; left ME: L-ME; left MI: L-MI).

**Figure 1:**
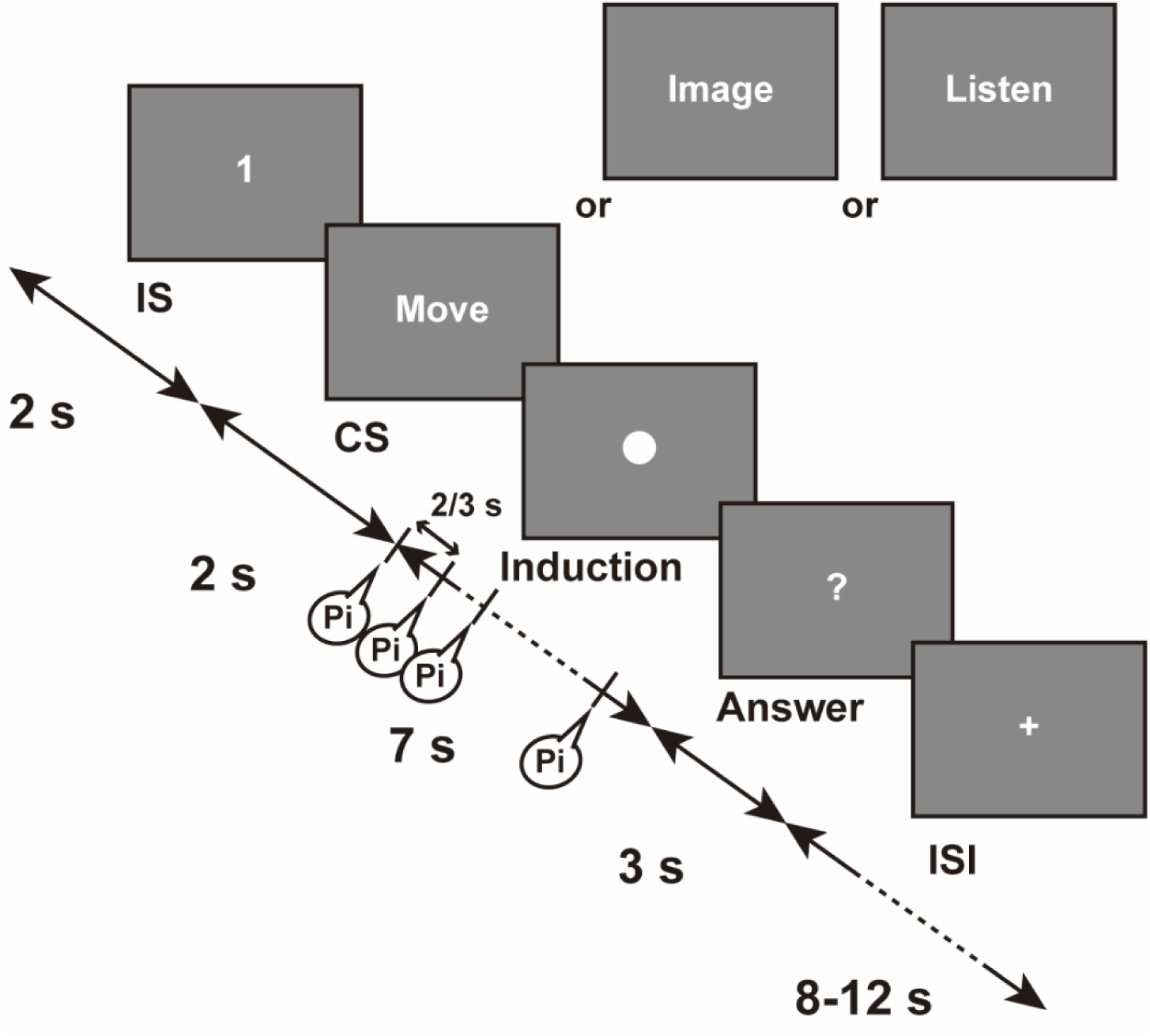
Design of the delayed sequential motor imagery and execution (dSMI) task. A participant was asked to remember the instruction stimuli (IS), which were presented for 2 s (1: pointing finger; 2: middle finger; 3: ring finger) and specified a segment of the cue stimuli (CS), which showed the following conditions: Move, Image, and Listen. Based on the CS, the participants sequentially tapped their fingers (Move: motor execution, ME), or imagined the tapping (Image: motor imagery, MI), or did nothing (Listen: passive listening, PL) during the induction period (IP).

### 2.3 MRI data acquisition

Images were acquired with a 3T MRI scanner, MAGNETOM Trio Tim (Siemens Medical Systems, Erlangen, Germany) installed in the Brain Activity Imaging Center in the ATR. High-resolution T1-weighted structural images were acquired for normalization to a standard brain for echo planar image (EPI) registration purposes (repetition time: TR = 2300 ms, echo time: TE = 2.98 ms, flip angle = 9°, inversion time: TI = 900 ms, matrix = 256 × 256, field of view = 256 mm, slice thickness = 1 mm, iso-voxel). Functional images were acquired with an EPI sequence (TR = 2000 ms, TE = 30 ms, flip angle = 80°, matrix = 64 × 64, field of view = 212 mm, slice thickness = 3.2 mm, gap: 0.8 mm, 33 slices, scan sequences: interleaved) of 536 volumes in the dSMI task for 17 min 52 s.

### 2.4 Data analysis

#### 2.4.1 Preprocessing of fMRI data

We followed a preprocessing protocol from a previous study (Ogawa et al., 2018). The data were processed using SPM12 (Wellcome Trust Centre for Neuroimaging). The first four volumes were discarded to allow for T1 equilibration. The remaining data were corrected for slice timing and realigned to the mean image of that sequence to compensate for head motion. Next, the structural image was co-registered to the mean functional image and segmented into three tissue classes in the MNI space. Using associated parameters, the functional images were normalized and resampled in a 2 × 2 × 2 mm grid. Finally, they were spatially smoothed using an isotropic Gaussian kernel of 8 mm full-width at half maximum.

#### 2.4.2 Regions of interest

We followed a previous study (Zabicki et al., 2017) which used sixteen brain regions and added bilateral dorsolateral prefrontal cortex (DLPFC) as control regions. Therefore, we defined eighteen ROIs (Table 1) as follows: frontal motor regions (M1, dPMC, vPMC), posterior parietal regions (SPL, IPL, IPS), and unrelated regions (primary auditory cortex: A1, fronto-marginal gyrus: FMG, dorsolateral prefrontal cortex: DLPFC). The anatomical masks were reconstructed from the Destrieux Atlas (Destrieux et al., 2010) in the Wake Forest University Pick Atlas (Maldjian et al., 2003). These masks were resliced to match the preprocessed EPIs with the same spatial resolution.

**Table 1.** A list of regional interests applying for the network analysis.

| # | ROI | Network | # | ROI | Network |
| --- | --- | --- | --- | --- | --- |
| 1 | L-M1 | Frontal, motor | 10 | R-M1 | Frontal, motor |
| 2 | L-dPMC | Frontal, motor | 11 | R-dPMC | Frontal, motor |
| 3 | L-vPMC | Frontal, motor | 12 | R-vPMC | Frontal, motor |
| 4 | L-SPL | Parietal | 13 | R-SPL | Parietal |
| 5 | L-IPL | Parietal | 14 | R-IPL | Parietal |
| 6 | L-IPS | Parietal | 15 | R-IPS | Parietal |
| 7 | L-A1 | Control, auditory | 16 | R-A1 | Control, auditory |
| 8 | L-FMG | Control, cognition | 17 | R-FMG | Control, cognition |
| 9 | L-DLPFC | Control, cognition | 18 | R-DLPFC | Control, cognition |

For each participant, we extracted fMRI time courses within each ROI. To remove several sources of spurious variance along with their temporal derivatives, linear regression was performed, including six motion parameters in addition to averaged signals over gray matter, white matter, and cerebrospinal fluid. Furthermore, to reduce spurious changes caused by head motion, the data were checked by a method that reduces motion-related artifacts. A band-pass filter (transmission range, 0.008−0.1 Hz) was applied to these sets of time courses prior to the following regression procedure. Eleven participants were excluded from the fMRI analysis because the participants did not perform all sessions of the experiment or had excessive head/body movements during the fMRI scans.

#### 2.4.3 DirectLiNGAM

Linear non-Gaussian acyclic causal model (LiNGAM: Shimizu et al., 2006) is a causal discovery model that utilizes higher-order distributional statistics to determine directed network connections. The assumptions of LiNGAM are as follows: 1) each node (variable) is linearly related to the other nodes, 2) its residual or external input has non-Gaussian distributions, and 3) the corresponding directed dependency graph is acyclic, i.e., the nodes have a natural ordering. Letting *x_i_* denote the *i*-th observation variable (*i =* 1*, …, n*), its linear association to the preceding/parent variable can be represented as

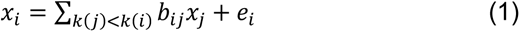

where *b_ij_* is the connection weight, quantifying the direct causal effect, and *k* (*i*) is the causal order for each variable. The external influences *e_i_* have non-Gaussian distributions, with non-zero variances, and are independent of each other. Rearranging equation (1) into a matrix form, we have

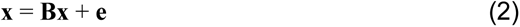

where **x** and **e** are the vectors of observed variables *x_i_* and external influences *e_i_*, respectively; **B** = (*b_ij_*) is an *n* × *n* connectivity matrix that can be permuted to a strictly lower triangular matrix (i.e. with all zeros in the main diagonals as well as the upper triangular part). Then, the identifiability of matrix **B** and the causal ordering/strength among the variables can be rigorously proven (Shimizu et al., 2006) based on the theory of independent component analysis (Hyvärinen et al., 2001).

We specifically used the DirectLiNGAM algorithm to estimate the strength of directed functional connectivity (i.e. nonzero entries of B) among brain regions for each of the MI/ME conditions. DirectLiNGAM requires no algorithmic parameters and is guaranteed to converge to a solution within a small number of steps if the data strictly follows the assumptions with sufficiently large samples, e.g., > 2,000 data points as suggested by previous studies (Smith et al., 2011; Xu et al., 2014). During the optimization process, the algorithm also prunes redundant edges and selects significant edges in an acyclic manner. This is important to concisely visualize the directed acyclic graph (DAG) with multiple ROIs and interpret differences across conditions. We used an open-source MATLAB implementation of DirectLiNGAM (Shimizu et al., 2011).

We first prepared a matrix of time courses of the fMRI signal that extracted six data sample epochs from the onset of the IP for each ROI (number of voxels × 720 samples; ME and MI conditions contained 30 trials; Fig. 2A). After normalizing the matrix by the mean and variance of the matrix, we conducted a principal component analysis for each ROI to decompose the loadings and scores (Fig. 2B). We corrected the sign of the first principal component (PC1) so that it has a positive correlation with the mean time course within the ROI. We collected the corrected PC1 across all ROIs and concatenated the matrix across all subjects for each condition (18 ROIs × 2,880 samples [6 samples × 30 trials × 16 subjects], Fig. 2C). We applied DirectLiNGAM for each condition, and estimated the **B** matrix, the nonzero entries of which identify the direct causal effects, i.e., directed functional connectivity, between two nodes, as well as the acyclic causal ordering of the nodes.

**Figure 2:**
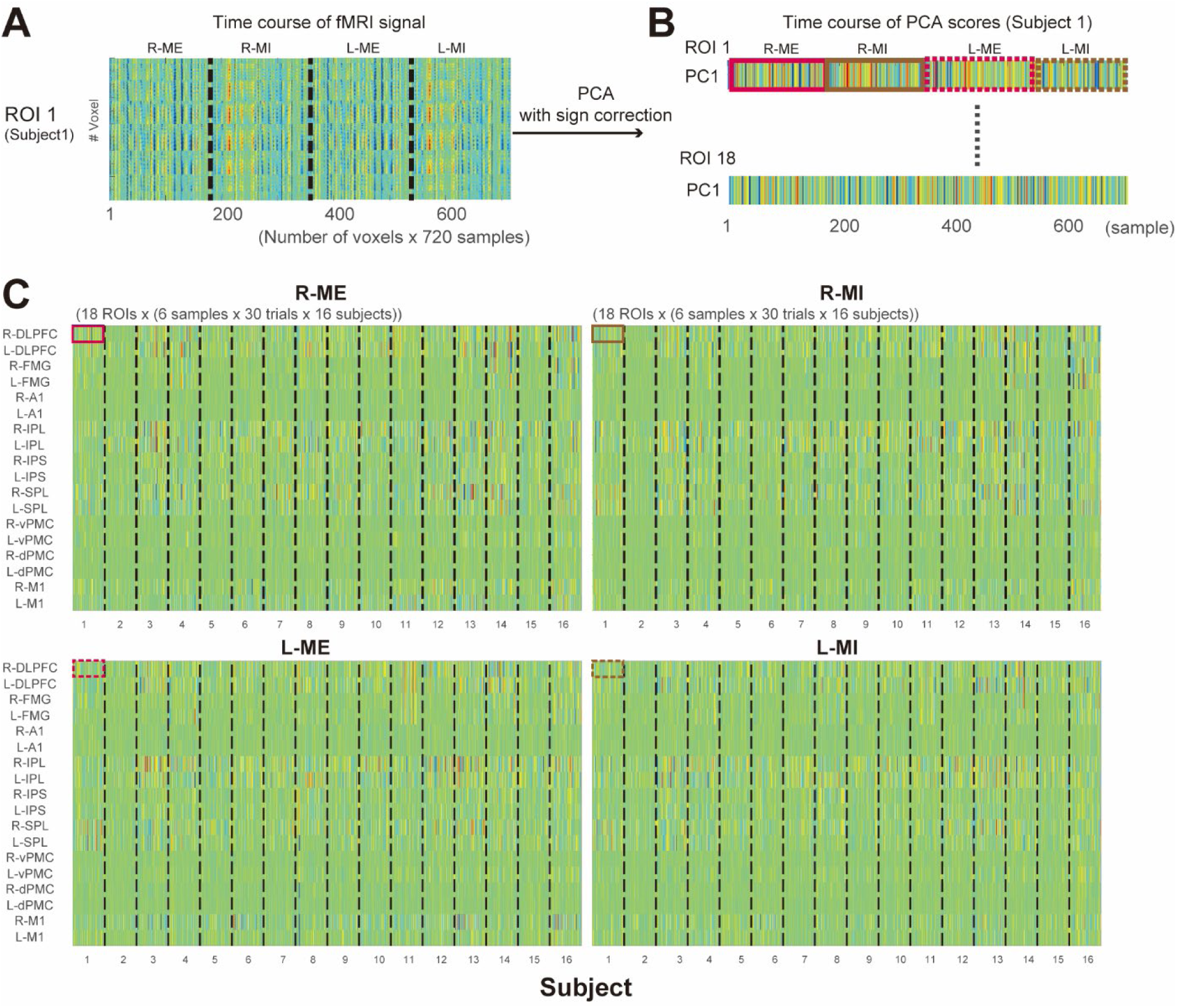
Preprocessing of input data matrix for DirectLiNGAM. (A) An example of a data matrix in single ROI. (B) Time courses of scores of the principal component analysis (PCA) after normalizing the matrix by mean and variance of the matrix for each ROI to decompose loadings and scores. (C) Input data matrix, x, for DirectLiNGAM algorithm. x was collected from only the first principal component (PC1) with sign correction using temporal correlation between the PC1 and a mean time course of the same ROI and concatenated across all subjects for each condition (ME, MI: 18 ROIs × 2,880 samples). L/R-ME: motor execution with the left/right hand; L/R-MI: motor imagery with the left/right hand; L/R-FMG: left/right frontomarginal gyrus; L/R-DLPFC: left/right dorsolateral prefrontal cortex; L/R-vPMC: left/right ventral premotor cortex; L/R-dPMC: left/right dorsal premotor cortex; L/R-M1: left/right primary motor cortex; L/R-SPL: left/right superior parietal lobule; L/R-IPS: left/right inferior parietal sulcus; L/R-IPL: left/right inferior parietal lobule; L/R-A1: left/right primary auditory cortex.

#### 2.4.4 Statistics of network strength

To evaluate the stability of directed functional connectivity across the subjects, we applied the bootstrapping method to estimate the lower and upper boundary of the strength of the connections with 100 repetitions following a method described by Xu et al. (2014). We assumed that the fMRI data, **x**, can be linearly represented by Eq. (2). To check the validity of the DirectLiNGAM approach, we confirmed non-Gaussianity of innovations, **e** = (**I**−**B**) **x**, of the estimated model by the Kolmogorov–Smirnov test (p < 0.05, “kstest.m”, in Statistics toolbox, MATLAB). We then estimated 100 models for each condition (R-ME, R-MI, L-ME, and L-MI) and applied the bootstrapped t-test for each condition (t-test, p < 0.001 corrected by the Bonferroni method). Additionally, to evaluate statistical difference of the direct effects between the two conditions, we applied the two-sample t-test (p < 0.001 corrected by Bonferroni method).

#### 2.4.5 General linear model

A first-level subject-wise analysis was performed using the general linear model (GLM) in SPM12. T-statistics were calculated for each subject using boxcar regressors, convoluted with a canonical hemodynamic response function without time and dispersion derivatives. The preprocessed individual functional images were high-pass filtered with a 128 s cutoff period to remove the effect of low-signal drift. Additionally, the six head movement parameters derived from the realignment procedure were used as regressors of no interest. fMRI data of each session were modeled with two regressors of interest (ME and MI) corresponding to the task blocks included in the session. All brain activity during the 7 s of the IP was included in the analysis. To identify the activated brain areas during the MI and ME conditions, contrast images were calculated including (i) R-ME > L-ME; (ii) L-ME > R-ME; (iii) R-MI > L-MI; (iv) L-MI > R-MI; (v) R-ME > R-MI; (vi) R-MI > R-ME; (vii) L-ME > L-MI; (viii) L-MI > L-ME. The contrast images were used as the input data in group-level random-effect analyses using one-sample *t*-tests. The threshold for statistical significance was set at an uncorrected p-value < 0.001 with a cluster-based family-wise-error correction of p-value < 0.05.

## 3. Results

### 3.1 Directed functional connectivity on fMRI data

Applying the DirectLiNGAM algorithm, we estimated the weight coefficient matrices, **B**, from data **x** for each condition (R-ME, R-MI, L-ME, L-MI). By shuffling the subjects’ order with 100 repetitions for each condition, we illustrated colormaps of the mean weights over repetitions, such that only significant ones were shown (Fig. 3A). According to the colormaps, we drew DAGs to visualize directed effects between two regions (Fig. 3B). To indicate graphs of the network, we counted number of edges a node has to or from other nodes as a measure of node importance in the network, (i.e., out-degree refers to the number of edges from the node to other nodes, and in-degree refers to the number of the edges from other nodes to the node) in Fig. 3CD.

**Figure 3:**
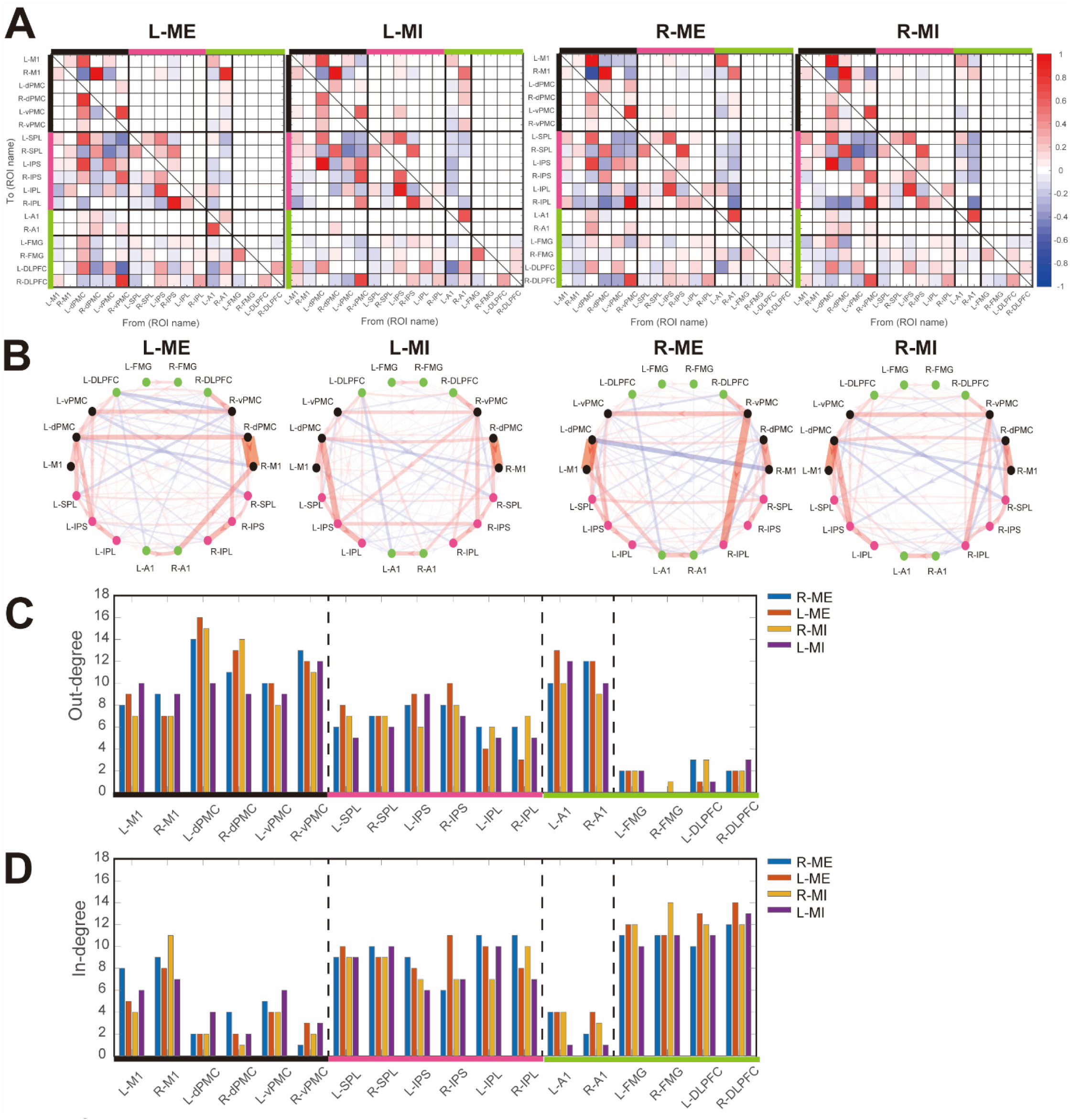
Direct acyclic graphs and matrices of directed functional connectivity estimated by DirectLiNGAM. The colormaps (A) and direct acyclic graphs (B) show mean strengths of directed functional connectivity, **B**, between two brain regions. Significant edges were connected between two ROIs (bootstrapping with 100 repetitions, thresholding p<0.001 corrected by Bonferroni method). ROIs were divided into three parts, frontal motor regions (black bars and dots in B), parietal regions (pink bars and dots) and control regions (green bars and dots) in A and B. The width of the edge in B shows the absolute value of the strength of the directed functional connectivity and an arrow of the edge is the direction. Positive weights are brightly colored and negative weights are in a blue colored. C and D: out-/in-degrees were counted with respect to significant functional directed connectivity for each ROI. Bars were colored by blue for R-ME, orange for L-ME, yellow for R-ME, and purple for L-MI.

First, brain regions in the bilateral d/vPMC showed high out-degrees and low in-degrees. This indicates that the bilateral d/vPMC strongly influences other regions, but direct effects from other regions are small (out-degree > in-degree, Fig. 3C and 3D). According to the colormaps and DAG, we determined the most dominant connection from dPMC to M1 contralateral to the hand (L-ME and L-MI: R-dPMC → R-M1; R-ME and R-MI: L-dPMC → L-MI; Fig. 3AB, Supp Table 1). We identified positive direct effects from the dPMC to M1 ipsilateral to the hand (R-ME and R-MI: R-dPMC → R-M1; L-ME and L-MI: L-dPMC → L-M1). As a bilateral interaction, we found positive direct effects (all conditions: L-vPMC → R-vPMC; L-ME, R-ME, L-MI: L-dPMC → R-dPMC). Additionally, we found strong negative direct effects from the L-dPMC to R-M1 in R-ME, but also weak negative effects in the other three conditions (all conditions: L-dPMC → R-M1). Most of positive direct effects in the same hemisphere showed symmetric structure; however, the interhemispheric directed functional connectivity with negative effects showed inhibitions in the target region and an asymmetric structure.

Second, we also found that brain regions in the bilateral A1 regions showed high out-degrees and low in-degrees (Fig. 3C and 3D). In the colormap of the B matrix and DAG, we found a significant positive effect, e.g., in the audio-motor network (L-ME and L-MI: R-A1 → R-M1; R-ME and R-MI: L-A1 → L-M1), and a weak negative connection to the parietal regions. We added A1 into our analysis as control regions based on a previous study (Zabicki et al., 2017), which was not expected motor processes associated with MI and ME. However, our results suggest that A1 may influence not only M1, but also parietal regions similar to d/vPMC during the dSMI task, because we used a sequential pitch to lead finger tapping smoothly for the participants.

Third, we found that in- and out-degree in the parietal regions were not obviously different compared to that in the frontal, auditory, and control regions. Connections in the frontoparietal network, in particular, from the d/vPMC to SPL/IPS/IPL were identified. We found direct effects from the frontal regions (d/vPMC) to parietal regions (SPL, IPL, IPL), for instance, L-ME (L-dPMC ◊ L-SPL, L-IPS; L-vPMC → L-IPS; R-vPMC → R-IPS), L-MI (L-dPMC → L-IPS; R-vPMC ◊ L-IPS, R-IPS, R-IPL), R-ME (R-vPMC → R-IPL; L-vPMC, L-dPMC ◊ L-IPS), and R-MI (L-dPMC → L-SPL, L-IPS; R-vPMC → R-IPL, R-IPS). Meanwhile, connectivity from the parietal regions to frontal regions was not selected by DirectLiNGAM. The direct effects from the frontal to parietal regions showed an asymmetric structure. In the parietal regions, the bilateral IPS seed regions seemed to influence neighboring regions such as the IPL and SPL across all conditions. Thus, to achieve MI and ME, direct effects may show interactions within the frontoparietal network and intraparietal network.

Regarding the usage of bilateral FMG and DLPFC as control regions, profiles of their in-/out-degrees were opposite to those of the d/vPMC and A1, such that the in-degrees of these regions were higher than the out-degrees. Additionally, directed connections from the FMG and DLPFC to frontal motor regions, the parietal regions, and A1 were not selected by DirectLiNGAM (Fig. 3A) across all conditions. This result indicates that brain activity in the FMG and DLPFC were not caused by the achievements of ME and MI.

### 3.2 Differences of direct effects due to modes and laterality

To identify the direct effects between the two conditions, we evaluated differences between two samples (A: L-ME vs. L-MI, B: R-ME vs. R-MI, C: L-ME vs. R-ME, and D: L-ME vs. L-MI in Fig. 4) with a two-sample t-test (p < 0.001 corrected by Bonferroni method). Each element showed a difference between two conditions of a direct effect which was displayed only if the difference was statistically significant. First, we found higher effects from the dPMC to M1 contralateral to the hand (green dotted boxes in Fig. 4C and 4D). Second, frontoparietal connectivity found L-dPMC → R-IPS and R-vPMC → R-IPS in L-ME, and R-dPMC → R-IPS and L-vPMC → R-IPS in R-ME (orange dotted boxes).

**Figure 4:**
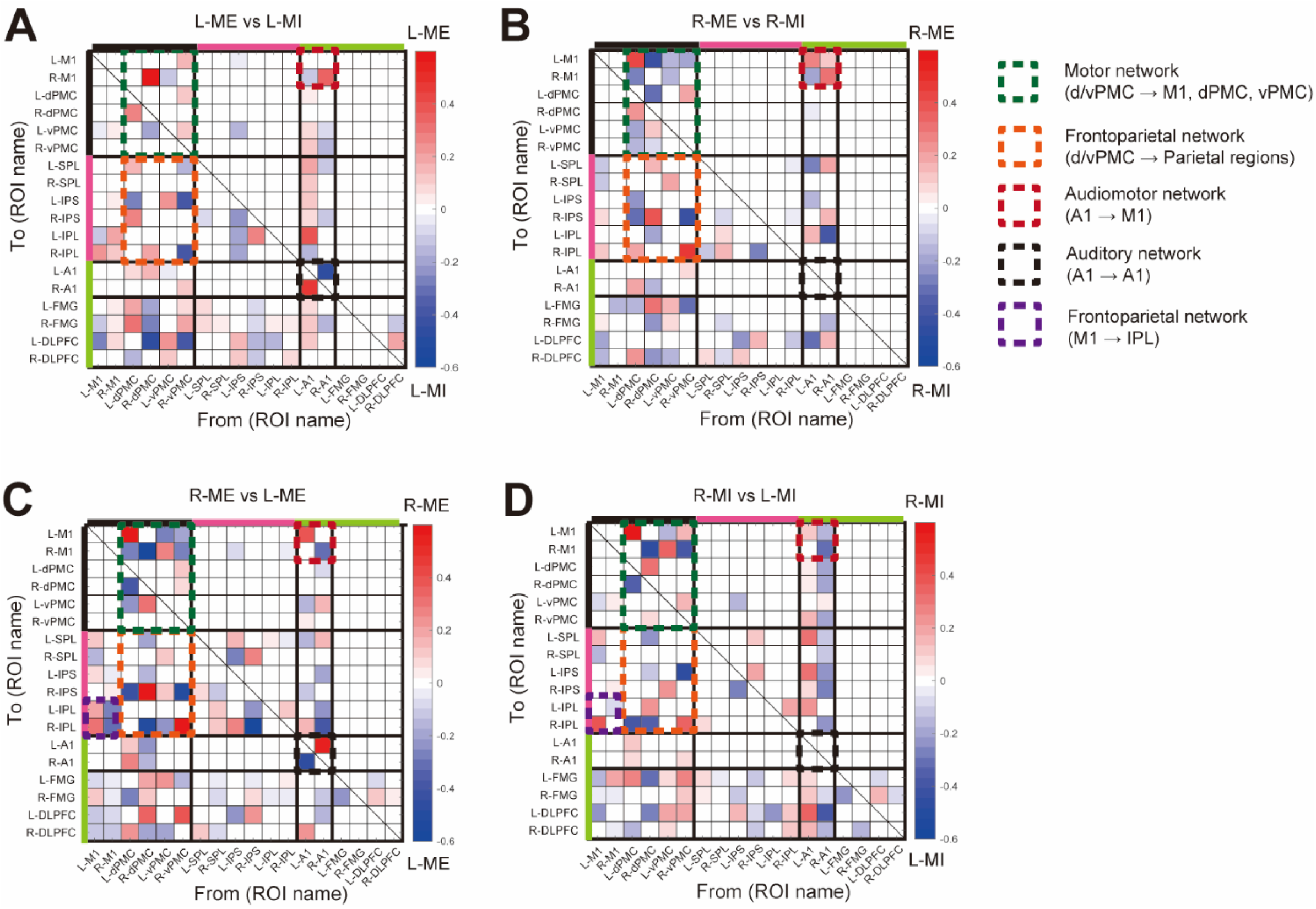
Directed graphs representing the differences between two conditions. From two samples estimated from bootstrapping, we illustrated significantly different edges with a two-sample t-test (p < 0.001 corrected by Bonferroni method). In (A) L-ME vs L-MI, (B) R-ME vs R-MI, it shows differences between ME and MI. The edges are colored by blue (ME) to red (MI). In (C) R-ME vs L-ME, (D) R-MI vs L-MI, it shows differences between R and L. The edges are colored by blue (ME) to red (MI). ROIs were divided into three parts, frontal motor regions (black bars), parietal regions (pink bars) and control regions (green bars) L/R-ME: motor execution with the left/right hand; L/R-MI: motor imagery with the left/right hand; L/R-FMG: left/right frontomarginal gyrus; L/R-DLPFC: left/right dorsolateral prefrontal cortex; L/R-vPMC: left/right ventral premotor cortex; L/R-dPMC: left/right dorsal premotor cortex; L/R-M1: left/right primary motor cortex; L/R-SPL: left/right superior parietal lobule; L/R-IPS: left/right inferior parietal sulcus; L/R-IPL: left/right inferior parietal lobule; L/R-A1: left/right primary auditory cortex.

Obvious differences in directed functional connectivity are represented in the edges dPMC → M1 and A1 → M1, contralateral to the hand (Fig. 4A and 4B). The direct effects, dPMC → M1, are more strongly affected in ME than in MI. This edge is essential to achieve motor control and related imagery. Second, the strength of the edges, A1 → M1, is also higher in ME and MI. It may represent a clear neural representation in both regions in the case of ME.

Comparing the laterality of ME (Fig. 4C) and MI (Fig. 4D), the edges dPMC → M1 contralateral to the hand clearly show differences in laterality. In comparison to ME, we identified edges L-M1 → R-IPL, R-dPMC → R-IPS larger in R-ME, and edges R-IPS → R-IPL in L-ME (Fig. 4C). In the case of MI, we identified edges L-M1 → L-SPL, R-M1 → L-FMG, R-IPL → R-DLPFC larger in R-MI, and edges, L-IPL → L-DLPFC, R-A1 → R-M1 larger in L-MI (Fig. 4D).

### 3.3 Results of general linear model of event related activity

To confirm brain activations between left and right ME/MI, we performed GLM analysis for comparisons of contrasts. First, we compared contrasts between R-ME and L-ME and summarized statistical results in Table 2. The brain regions that emerged from a contrast of R-ME > L-ME included the postcentral gyrus (M1), insula including the Rolandic operculum (RO) and Heschl gyrus (A1), putamen, SMA, and thalamus in the left hemisphere contralateral to the moving hand. In the opposite contrast, L-ME > R-ME, we identified clusters in the precentral gyrus, thalamus, putamen in the right hemisphere contralateral to the moving hand. In the case of MI, we found similar tendencies (R-MI > L-MI: L postcentral gyrus [L-M1], L insula; L-MI > R-MI: R postcentral gyrus [L-M1]). These results were consistent with previous findings that brain activity in the contralateral hemisphere is predominantly higher than that in the ipsilateral hemisphere. Therefore, the clusters were symmetrically colocalized in comparisons between lateralities (Hanakawa et al., 2008).

**Table 2.**
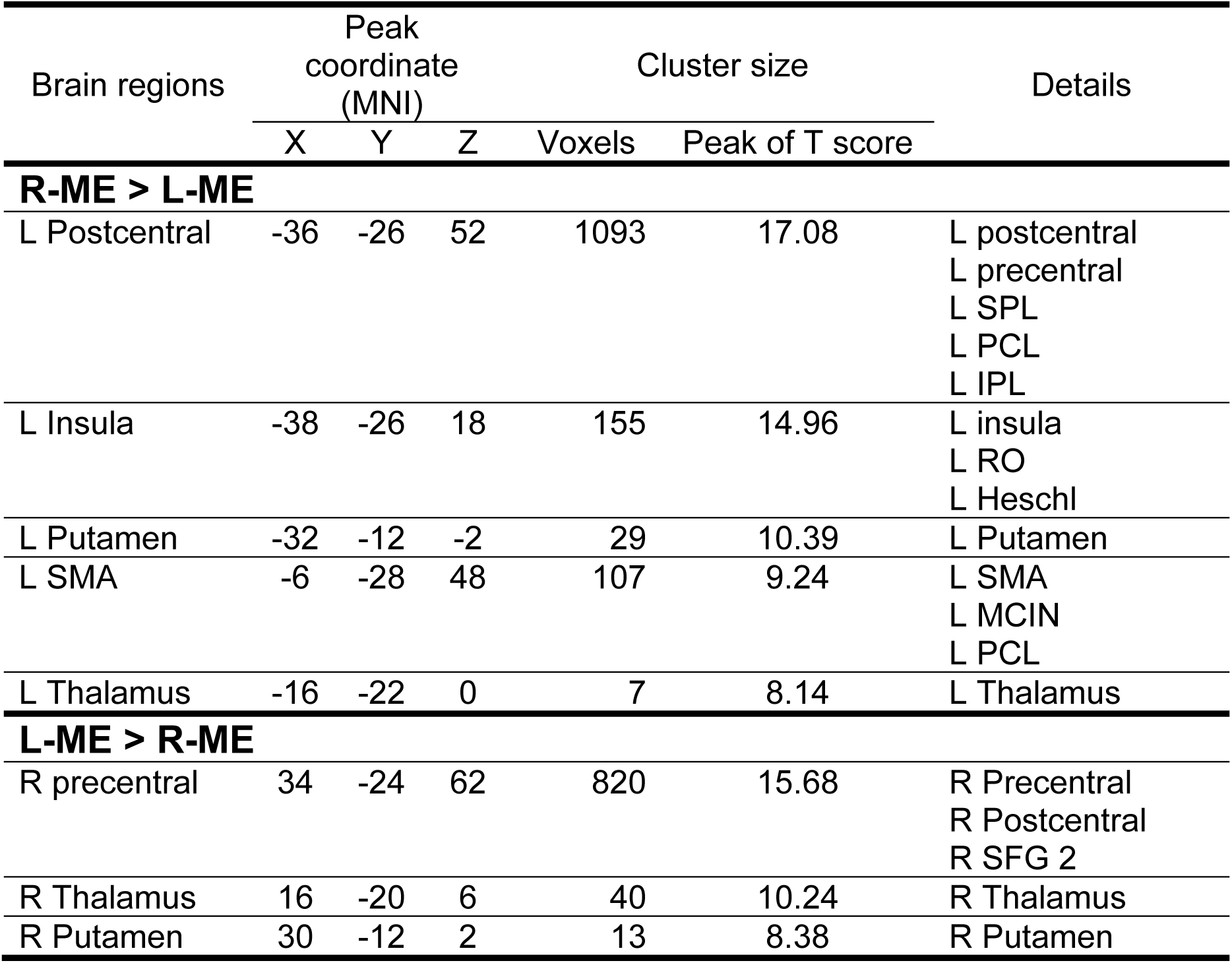
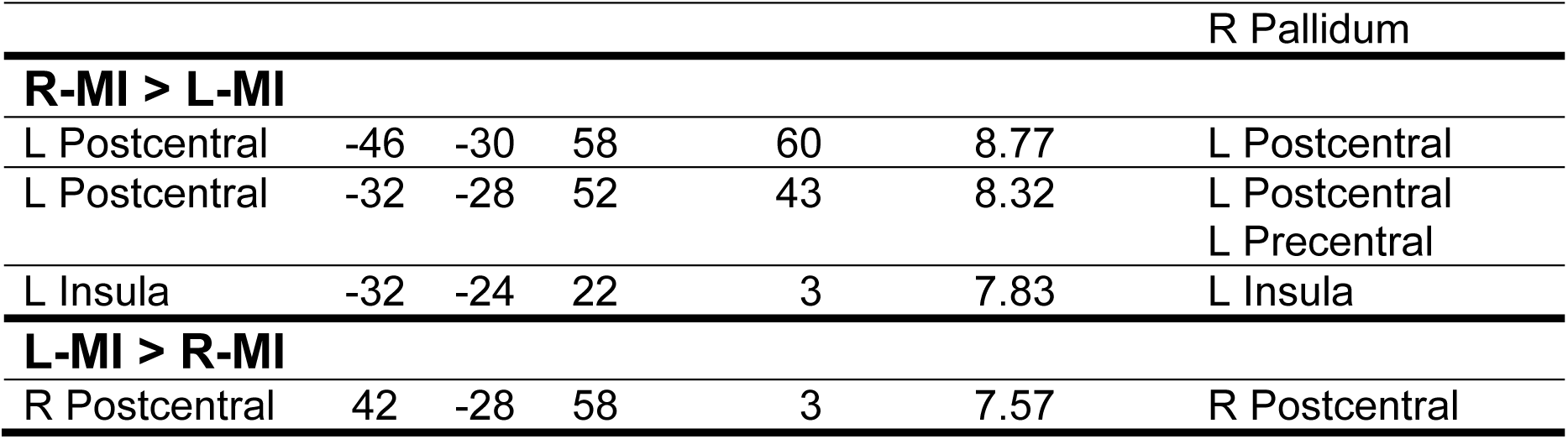
Results for the contrasts between right and left hands [R-ME > L-ME], [L-ME > M-ME], [R-MI > L-MI], and [L-MI > R-MI]. MNI coordinate is corresponding to the peak voxels within each cluster. Clusters were set at a threshold of p < 0.001 and cluster level family-wise-error at p < 0.05. Note: Brain regions were labelled by the AAL toolbox in the SPM extension. SPL: Superior parietal lobule; PCL: Paracentral lobule; IPL: Inferior parietal lobule; RO: Rolandic operculum; SMA: Supplementary motor area; MCIN: Middle cingulate gyrus, middle part; SFG: Superior frontal gyrus.

Next, we compared contrasts between ME and MI such as modes for each hand (Table 3). The brain regions that emerged from a contrast of R-ME > R-MI included the left postcentral gyrus (L-M1), which means that L-M1 is more activated at R-ME than R-MI. Similarly, we identified clusters with a contrast of L-ME > L-MI including those in the right precentral (R-M1) and right thalamus. On the other hand, opposite contrasts (R-MI > R-ME, L-MI > L-ME) showed clusters in the prefrontal (L inferior frontal gyrus, triangle part: IFG Tri, L middle frontal gyrus: MFG, R MFG), parietal (L IPL, L SPL), and occipital (L superior occipital gyrus: SOG, L middle occipital gyrus: MOG) regions. Brain activity corresponding to ME in M1 was higher than that of MI, but brain activity corresponding MI was lateralized in the left hemisphere.

**Table 3.** Results for the contrasts between ME and MI [R-ME > R-MI], [R-MI > R-ME], [L-ME > L-MI], and [L-MI > L-ME]. MNI coordinate is corresponding to the peak voxels within each cluster. Clusters were set at a threshold of p < 0.001 and cluster level family-wise-error at p < 0.05. Note: Brain regions were labelled by the AAL toolbox in the SPM extension. IFG: Inferior frontal gyrus; IFG Tri: IFG triangular; SFG: Superior frontal gyrus; MFG: Middle frontal gyrus; IFG Oper: IFG, opercular; SMA: Supplementary motor area; IPL: Inferior parietal lobule; SMG: Supramarginal gyrus; SPL: Superior parietal lobule; MTG: middle temporal gyrus; SOG: Superior occipital gyrus; MOG: Middle occipital gyrus; PCL: Paracentral lobule.

| Brain regions | Peak coordinate (MNI) |  |  | Cluster size |  | Details |
| --- | --- | --- | --- | --- | --- | --- |
|  | X | Y | Z | Voxels | Peak of T score |  |
| R-ME > R-MI |  |  |  |  |  |  |
| L Postcentral | -38 | -26 | 54 | 219 | 9.49 | L Postcentral<br>L Precentral |
| R-MI > R-ME |  |  |  |  |  |  |
| L IFG Tri | -36 | 10 | 18 | 3254 | 6.43 | L IFG tri<br>L MFG 2<br>L Precentral<br>L IFG Oper<br>L SFG 2<br>L SMA<br>L Postcentral<br>L Insula |
| L IPL | -56 | -44 | 50 | 1143 | 6.26 | L IPL<br>L SMG<br>L Angular<br>L SPL<br>L MTG |
| L SOG,<br>L SPL | -10 | -82 | 44 | 453 | 5.38 | L SOG<br>L SPL<br>L MOG<br>L Precuneus<br>L Cuneus<br>L IPL |
| **L-ME > L-MI** |  |  |  |  |  |  |
| R Precentral | 34 | -22 | 68 | 218 | 10.22 | R Precentral<br>R Postcentral |
| R Thalamus | 20 | -22 | 10 | 24 | 8.73 | R Thalamus |
| **L-MI > L-ME** |  |  |  |  |  |  |
| L MFG 2 | -34 | 60 | 16 | 2544 | 7.36 | L MFG 2<br>L IFG Tri<br>L SFG 2<br>L IFG Oper<br>L IFG Orb 2<br>L Precentral<br>L Insula |
| L MFG 2 | -30 | 4 | 60 | 234 | 6.14 | L MFG 2<br>L SFG 2<br>L Precentral |
| L IPL | -6 | -74 | 32 | 1902 | 6.06 | L IPL<br>L SPL<br>L Precuneus<br>L Cuneus<br>L SOG<br>L MOG<br>R PCL<br>L Angular |
| R MOG | 36 | -80 | 34 | 210 | 5.92 | R MOG<br>R SOG<br>R SPL<br>R Cuneus<br>R Precuneus<br>R Angular |
| R MFG 2 | 40 | 42 | 34 | 207 | 5.52 | R MFG 2<br>R IFG Tri<br>R IFG Oper |

## 4. Discussion

In this study, we demonstrated the asymmetric representation of the direct functional connectivity during ME and MI. By applying DirectLiNGAM and GLM to the fMRI data, we characterized functional asymmetry in the d/vPMC and A1, parietal regions, and prefrontal control regions. We found higher direct functional connectivity from the contralateral dPMC to M1 compared to that from the ipsilateral dPMC to M1, as well as activations in M1. Regarding the following of finger tapping with beep sounds, A1 showed high out-degrees that affected M1 predominantly, as well as other regions. Regarding the parietal regions, strong connections were observed mainly from the d/vPMC, as well as intra-connections. The frontal control regions were affected by other regions, but not vice versa. However, higher activations during MI were distributed in the left hemisphere than ME in both cases of the right and left hand, such that MI showed functional asymmetry. Here, we would like to discuss the network structure with directed functional connectivity and its brain activations across hands and modes.

### 4.1 Functional roles of d/vPMC evidenced by activation patterns and causal discovery during ME and MI

Our results of the GLM in Tables 2 and 3 showed that the brain activity is generally contralateral to the movement/imagery side, and that brain activity in M1 during ME was higher than that during MI. They were consistent with findings from previous neuroimaging studies (Kawashima et al., 1993; Hanakawa et al., 2003; Gao et al., 2011). From a viewpoint of the causal discovery analysis, strong positive direct effects, dPMC → M1, in the same hemisphere contralateral to the tapping hand during MI and ME were selected (Fig. 3). Previously, positive directed functional connectivity has been found from the dPMC/SMA to M1 (CGCM: Gao et al., 2011, Wang et al., 2019; DCM: Kasess et al., 2008), therefore, the positive directed functional connectivity shows active inference from a source region to a target region. Gao et al. demonstrated that the out-degrees of the dPMC contralateral to the performing hand were higher than the in-degrees regardless of ME/MI tasks with right/left hand. Our results also showed that the out-degrees of dPMC were higher than the in-degrees. This indicates that dPMC is one of the generators of neural representation associated with ME/MI, not just in M1 but also in the parietal regions.

We also found positive effects, dPMC → M1, ipsilateral to hand (Supp. Table 1). Our task requires the participants to tap or imagine tapping with one hand; therefore, we did not expect to identify the ipsilateral connections between the dPMC and M1 in this analysis. Comparisons the right and left hemispheres, the contralateral connections were higher than the ipsilateral ones in both ME and MI. In a previous study by Gao et al. (2011), functional connectivity was estimated by CGCM with ME/MI of the right/left hand similar to our task; however, they did not include ipsilateral M1 into the model. Wang et al. (2019) also estimated functional connectivity of five ROIs by CGCM during ME and MI of young and old subjects, and they found bilateral connections between the PMC and M1. We successfully identified the ipsilateral connections, dPMC → M1, despite the high dimensional network compared to the previous studies. It suggests that neural representations in the ipsilateral pathway may partly help to achieve ME and MI.

### 4.2 Audio-motor network during ME and MI

Regarding the audio-motor network, we found i) high out-degree and low in-degree, ii) positive effects, A1 → M1 bilaterally, and iii) negative effect, A1 → parietal regions, in particular, connections within the same hemisphere. We did not expect to find connections related to A1 when we designed this study because we assumed that A1 was a control region. The decoding accuracies of modality (ME vs. MI) in A1 obtained by multi-voxel pattern analysis were significant in a study by Zabicki et al. (2017). It suggests that the voxel pattern in A1 contains information that can distinguish between ME and MI. Lima et al. (2016) reviewed the roles of the SMA and pre-SMA in auditory processing and auditory imagery and concluded that activating sound-related motor representation in the SMA and pre-SMA might facilitate behavioral responses to sounds. On the other hand, Morillon et al. (2015) mentioned that connections between auditory and premotor/motor areas had not been established and were either weak or absent in primates. Our results indicate that there are direct effects on the audio-motor pathway, to M1 bilaterally and positively, and to the parietal regions negatively in the same hemisphere. In our task, the participants were instructed to follow the beep sounds to tap their finger or imagine finger tapping, without the inclusion of spatial information in sounds. Periodic sounds associated with motor control may facilitate the preparation of the next finger movement following the sound sequence and inhibit spatial audio-motor processes according to negative effects from A1 to the parietal regions observed in L-MI and R-MI (Fig. 3A).

### 4.3 Functional asymmetry of the parietal regions and left DLPFC

Regarding networks associated with the parietal region, we found functional asymmetry, frontoparietal connections (negative effects: bilateral-vPMC → bilateral SPL; positive/negative effects: bilateral dPMC → bilateral SPL, L-IPS), common connections within the parietal regions (L-SPL → L-IPL; R-SPL ◊ L-IPL; L-IPL → L-SPL, R-IPS, L-IPL; R-IPS → R-SPL, R-IPL), and that the left parietal regions were highly activated in MI compared to ME. In a previous study that used CGCM, bidirectional directed functional connectivity between the dPMC and SPL/IPL in the same hemisphere was identified from fMRI data during ME and MI (Gao et al., 2011). In their results, bidirectional directed functional connectivity represents the forward and backward interactions between the dPMC and SPL during the visual-cued motor task. It suggests that SPL may be associated with the process of somatosensory modality and visual information and its integration. There are functional similarities and differences in PMC across the dorsal-ventral axis, such as the prediction of sequential finger movement (Nambu et al., 2015), representation of chuck of sequential finger movement (Yokoi & Diedrichsen, 2019), encoding of spatial location (Gallivan et al., 2011), and linguistic motor planning of fingers (Hanakawa et al., 2005). On the other hand, a specific role of vPMC is to encode target locations depending on head/eye-centered coordinates and arm/limb-centered coordinates (Gallivan et al., 2011). In a study by Hanakawa et al. (2005), neural representations in vPMC mediate the mapping of linguistic stimulus onto the motor planning of fingers. Due to motor and non-motor sequences, activations in the vPMC are enhanced during ME and MI tasks. It suggests that functions in the vPMC are associated with encoding effectors and spatial coordination.

Additionally, regarding the comparisons of brain activity, the left parietal regions had significantly greater activation during MI compared to ME (Table 2 and 3). L-IPS has positive effects to the L-SPL, R-IPS, and L-IPL across all conditions. Interestingly, these results indicate the functional asymmetry of the parietal regions. This is inconsistent with the results of Gao et al., such as the directed functional connection between frontoparietal regions (Gao et al., 2011). The parietal regions are important to encode spatial processing of target or effector-specific coordinates, such that neural representation of gestures/actions with the hand/foot have been predicted by classification methods (three right-hand actions: aiming, extension-flexion, and squeezing: Pilgramm et al., 2016; three right-hand actions and their imagery: Zabicki et al., 2017; touching left/right and gazing left/right: Gallivan et al., 2011). Our result was comparable to previous findings even though our task involves the modulation of the laterality of the hands (right/left) and modes (MI/ME) with just finger movement.

Regarding the left DLPFC, we identified clusters satisfying the condition MI > ME (Table 3). In early neuroimaging studies, functional asymmetry was found in the M1 and left prefrontal regions (Kawashima et al., 1993), precuneus, and middle frontal gyrus (Hanakawa et al., 2003). The results of DirectLiNGAM showed that the DLPFC had high in-degree and low out-degree with weak connections, indicating that it may collect sensory information. In particular, MI induced not only motor-related activity in the frontoparietal network, but also activity in the prefrontal regions which integrates information from the body and the environment and participates in higher-order motor control (van der Meulen et al., 2014).

### 4.4 Limitation of DirectLiNGAM

The DirectLiNGAM algorithm is a suitable approach to explore the causal relationships between multiple nodes (Shimizu et al., 2011). The conventional methods, such as DCM or GCM, requires large computational costs in treating four or more time series. One main limitation is that if brain regions A and B are both driven by region C but with a different lag, an effective connectivity will be shown an edge between A and B in the DCM. To address this problem, the GCM method has been proposed and applied to field potential data from macaque monkeys when performing a go/no-go visual pattern discrimination task (Zhang et al., 2008) and human fMRI data during a face matching task (Wang et al., 2017). Afterwards, GCM has also improved to enable its application to more than the data of three nodes employing a conditional variable similar to the partial correlation analysis (Wang et al., 2019).

Contrastingly, the DirectLiNGAM algorithm is identifiable by a multivariate dataset of more than three variables. The advantages of this algorithm are as follows: i) a full and unique causal structure can be identified, provided all assumptions hold and the sample size of the observations is sufficiently large, ii) no prior knowledge of the network structure is needed, and iii) compared to DCM, which may involve large numbers of conditional independent tests, the computational complexity of LiNGAM is much lower (Smith et al., 2011). However, DirectLiNGAM is assumed to be DAG and to prune weakly contributed edges with sparseness; therefore, there is a possibility of missing bidirectional directed functional connectivity. In this study, comparisons of a cyclic model or a LiNGAM model with hidden variables/confounders were out of scope. It is necessary to evaluate how the different bidirectional effects or latent confounders are represented in GCM and DirectLiNGAM in future studies.

## 5. Conclusions

By applying the causal discovery approach to the fMRI data, we demonstrated that DAG represented the frontoparietal motor network underlying ME and MI. To support the estimation of the directed functional connectivity, we also confirmed spatial brain activity with GLM during ME and MI. We developed a dSMI task based on that described by Hanakawa et al. (2008) to acquire enough fMRI data samples under the ME and MI conditions. We found predominant directed functional connectivity, d/vPMC → M1, which is a core of the motor control. Additionally, we identified audio-motor connections from A1. The parietal regions and DLPFC showed functional asymmetry during ME and MI. The parietal regions may not only receive movement related information from d/vPMC but may also communicate with neighboring regions. MI can cause higher cognitive load than ME. Our approach has the limitations of representing bidirectional interactions and the assumption of independence for each variable. It is necessary to compare several methods of the representation of the causalities of multiple brain regions. Our findings might help to understand large-scale interactions across bilateral frontoparietal networks during ME and MI.

## Acknowledgements

This study was supported by JSPS KAKENHI Grant Number JP18H05395, JP18KK0284, JP21K12620, JP21K12055, JP21H03516, the ImPACT Program of the Council for Science, Technology and Innovation (Cabinet Office, Government of Japan), a project, JPNP20006, commissioned by the New Energy and Industrial Technology Development Organization (NEDO), CREST, JST, a contract entitled “Novel and innovative R&D making use of brain structures” with the Ministry of Internal Affairs and Communications, Japan, and the MIC/SCOPE/ #192107002. We thank Manabu Shikauchi and Shigeyuki Ikeda for supporting the MRI experiments, Shohei Shimizu and Aapo Hyvärinen for advising the statistical interpretation of LiNGAM analysis.

## Declaration of interest

None

## Author contributions

Takeshi Ogawa designed this study and performed data acquisition. Takeshi Ogawa and Hideki Shimobayashi analyzed the data. Takeshi Ogawa, Hideki Shimobayashi, Jun-ichiro Hirayama, and Motoaki Kawanabe discussed data analysis and results. Takeshi Ogawa and Jun-ichiro Hirayama wrote the original draft.

## Data and code availability

The data that support the findings of this study are available from the corresponding author, Takeshi Ogawa, upon reasonable request.

## Notes

### Competing Interest Statement

The authors have declared no competing interest.

### Summary of Updates

Section on Results and Discussion to clarify; Figure 3 and 4 revised; Supplemental files updated.

