## Supplemental Table 1 for "Asymmetric directed functional connectivity within the frontoparietal motor network during motor imagery and execution"

### Supplemental materials

#### Supp. Table.1

**Comparisons of direct effects from the dPMC to M1 for each condition and each difference**

| | **L-ME** | **L-MI** | **R-ME** | **R-MI** |
|---|---|---|---|---|
| L-dPMC → L-M1 | + (ipsi) | + (ipsi) | + (contra) | + (contra) |
| L-dPMC → R-M1 | - | - | - | - |
| R-dPMC → L-M1 | Nan | Nan | - | + |
| R-dPMC → R-M1 | + (contra) | + (contra) | + (ipsi) | + (ipsi) |
| | **L-ME vs L-MI** | **R-ME vs R-MI** | **R-ME vs L-ME** | **R-MI vs L-MI** |
| L-dPMC → L-M1 | Nan | R-ME (ipsi) | R-ME (contra) | R-MI (contra) |
| L-dPMC → R-M1 | Nan | R-MI | L-ME | Nan |
| R-dPMC → L-M1 | Nan | R-MI | Nan | Nan |
| R-dPMC → R-M1 | L-ME (ipsi) | Nan | L-ME (contra) | L-MI (contra) |
